# The human DEAD-box protein DDX3X regulates host and viral mRNA translation during Sendai Virus infection

**DOI:** 10.64898/2026.03.08.707086

**Authors:** Cathal S. Ryan, Dimitrios G. Anastasakis, Ahsan H. Polash, Elizabeth Sitko, Markus Hafner, Martina Schröder

## Abstract

DDX3X is a multifunctional DEAD-box RNA helicase with important roles in translation initiation and antiviral innate immune signaling, yet it is currently unknown whether viral infection affects its interactions with host RNAs. Here, we define the transcriptome-wide binding landscape of endogenous DDX3X in Sendai virus-infected human cells using PAR-CLIP. We show that DDX3X maintains its preference for GC-rich, highly structured 5′UTR regions during infection, but acquires a distinct set of infection-induced targets, including *IFNB1* and multiple interferon-stimulated genes. We demonstrate that DDX3X directly binds the *IFNB1* 5′UTR and promotes its translation, establishing a previously unrecognized post-transcriptional mechanism contributing to DDX3X-dependent IFN-β production. We also evaluated DDX3X’s binding to SeV RNAs and concluded that DDX3X is likely not actively recruited by SeV or has a significant effect on its viral life cycle. Our findings add a novel dimension to DDX3X’s involvement in anti-viral immunity with implications for further therapeutic development of DDX3X inhibitors.

## INTRODUCTION

The human DEAD-box protein 3 X (DDX3X) is an RNA helicase involved in remodeling local secondary structures in RNAs in an ATP-dependent manner (1, 2). One of DDX3X’s main functions is the regulation of mRNA translation initiation, where it likely resolves structures in 5’ untranslated regions (UTRs) that may otherwise inhibit assembly or scanning of the ribosomal pre-initiation complex (3–6). DDX3X interacts with several core translation machinery components such as eukaryotic initiation factors (eIFs), ribosomal proteins, rRNAs, and several other RNA-binding proteins (RBPs) and is recruited to stress granules under cellular stress conditions (Ryan and Schröder, 2022; Lennox et al., 2020; Valentin-Vega et al., 2016; Vellky et al., 2020). Although several studies support a role for DDX3X in regulating mRNAs with structured 5’UTRs, the underlying determinants of DDX3X target selectivity remain incompletely understood (7, 11).

DDX3X has been found to be an essential host factor bound and recruited by a wide variety of viruses, ranging from HIV-1 (4, 5, 12) to SARS-CoV-2 (13). Because of this widespread exploitation of DDX3X helicase activity to support viral translation and/or genome replication, small-molecule inhibitors of DDX3X’s enzymatic activity are being explored for development as potential broad-spectrum antiviral drugs (14). However, this is complicated by the fact that DDX3X also enhances anti-viral innate immune signaling through an unconventional role as an adaptor molecule that does not require its enzymatic activity. DDX3X enhances activation of the kinase IKKε and bridges the interaction with its substrate, the transcription factor IRF3, which is required for transcriptional upregulation of the anti-viral cytokine *IFNB* downstream of the pattern recognition receptor RIG-I (15–17). The co-existence of these pro- and anti-viral functions implies that DDX3X’s role in viral infections is highly context-dependent and further characterization of its behavior in virus-infected cells will be necessary to inform its potential as an anti-viral drug target.

Despite well-documented interactions with various viral RNAs (7) it remains unknown how infection alters DDX3X’s interactions with the host cell transcriptome and whether this contributes directly to anti-viral immunity. Because host and viral RNAs compete for the cellular translation machinery and this can be decisive for infection outcome, it is important to understand infection-induced changes to DDX3X’s RNA target pool. Here, we use PAR-CLIP (18) to systematically map DDX3X RNA binding sites in uninfected and Sendai Virus (SeV)-infected cells. We show for the first time that DDX3X binds many infection-induced host mRNAs and that it can post-transcriptionally regulate expression of the key anti-viral cytokine IFNB, adding another dimension to DDX3X’s anti-viral functions during infection.

## MATERIALS AND METHODS

### Cell Culture and Virus Infection

HEK293T cells were grown in DMEM (Gibco, 21969035) supplemented with 10% FBS and Penicillin (100 U/mL) and Streptomycin (100 µg/mL). Cells were tested for Mycoplasma infection using Mycostrip® tests (Invivogen) according to the manufacturer’s instructions. Knockdown experiments used stable HEK293T cell lines containing doxycycline-inducible pTRIPZ constructs expressing either a short hairpin RNA (shRNA) targeting DDX3X (shDDX3X, V2THS_228965, Dharmacon/Revvity) or a non-silencing control shRNA (NSC, #RSH4743, Dharmacon/Revvity), which were previously described (Gu et al., 2017). Knockdown was induced for 72 hours at time of cell seeding via addition of 2μg/mL doxycycline. For SeV infections, cells were inoculated at a concentration of 80 Hemagglutinating-units/mL of Cantell-strain SeV purchased from Charles River Laboratories.

### RNAseq

For RNAseq, cells were seeded in 15 cm^2^ dishes at an initial concentration of 1 * 10^6^ cells/plate. When harvesting cells, media was aspirated and plates were washed with 5ml ice-cold PBS before 400ul of extraction buffer (20 mM Tris pH 7.4, 150 mM NaCl, 4 mM MgCl_2_, 1 mM DTT, 100 μg/ml Cycloheximide, 1% NP40, 25 U/ml Turbo™ DNase I (Invitrogen, AM2238)) was added. Cells were scrape-collected, left for 10min on ice, then aspirated at least 6 times through a 27G needle. Lysates were centrifuged for 10 min at 20,000 x g, 4°C. Total RNA was extracted in 1 mL Trizol (Invitrogen, 15596026) following the manufacturer’s instructions. RNAseq library preparation was performed using the NEBNext® Ultra™ II DNA Library Prep Kit (NEB, #E7645S/L) with rRNA depletion kit (#E6310) following the manufacturer’s instructions. Libraries were sequenced using an Illumina HiSeq (Illumina).

### PAR-CLIP

PAR-CLIP was performed as per Anastasakis *et al.* (2020). Briefly, for each sample, eight 15 cm dishes were seeded at an initial concentration of 5 * 10^6^ cells per plate. 16 hrs before harvesting, 100 µM 4-thiouridine was added to the plates. Medium was removed from plates, followed by washing with 10 ml PBS before crosslinking with 365 nm UV, 500 mJ/cm^2^. Cells were harvested, centrifuged at 300 x g for 10 min at 4°C, and frozen at -80°C until further processing.

Cell pellets were resuspended in 1ml RIPA buffer (20 mM Tris, pH 7.5, 150 mM NaCl, 0.1 % SDS, 0.15 % sodium deoxycholate, 1 % NP40, 0.5 mM DTT, protease inhibitor cocktail (Roche, 4693159001)) for 10 min, followed by three rounds of sonication for 30 seconds in between chilling for 30 seconds on ice. Lysates were then diluted in 4 ml of immunoprecipitation (IP) buffer (20 mM Tris, pH 7.5, 150 mM NaCl, 2 mM EDTA, 1% (v/v) NP40) before IP with protein G magnetic beads (Invitrogen, 10003D) pre-coupled with anti-DDX3X (Santa Cruz Biotechnology, 2253C5A) for 3 hours at room temperature (RT). After washing, beads were incubated with 0.0825 U/µl RNaseI (Thermo Fisher Scientific, AM2294) in IP buffer for 10 min at 22°C with shaking, washed, and ligated to a fluorescent 3’ RNA adapter (Table S1) overnight at 4°C with shaking. Samples were then phosphorylated with T4 PNK Kinase (NEB, M0201S) for 30 min at 37°C with shaking and then heated to 95°C for 5 min in 2x NuPAGE LDS sample buffer + DTT (Thermo Fisher Scientific, NP0008, NP0009), then loaded and resolved on Criterion™ XT Bis-Tris 4–12% gels (Bio Rad, #3450123). Gels were imaged for fluorescence, and bands corresponding to the apparent molecular weight of DDX3X plus 20kDa (∼100 kDa) were then excised from the gel. Protein was digested within the excised gel segments via treatment with Proteinase K (Millipore Sigma, 3115879001). RNA was recovered via phenol-chloroform extraction and precipitation with 10 mg/mL GlycoBlue (Thermo Fisher Scientific, AM9515) in 1 mL isopropanol. Recovered RNA was ligated to a 5’ RNA adapter (Table S1) using T4 RNA ligase (Thermo Fisher Scientific, EL0021) before reverse transcription, PCR amplification, and size selection for 78-100 nt on a 3% agarose Pippin Prep gel (Sage) as previously described (18). After a final PCR, samples were sequenced using an Illumina HiSeq (Illumina).

### NGS data processing

For PAR-CLIP, reads were processed, and peak calling was performed using ‘pcliptools-align’ from PCLIPtools (v.0.7.3) as in (19). Briefly, PCR duplicates were collapsed, and adapters removed using Cutadapt (v.4.5) (20). Reads were then aligned to a reference genome using STAR (v0.7.9a) (21). Alignments containing template mismatches were then extracted using Samtools (v1.2) (22) and Bedtools (v2.31.1) (23), and a Poisson model was computed based on the rate of mismatches. For contiguous sets of aligned reads that exceeded a depth threshold, peak calling was performed by determining whether a higher-than-expected number of T-to-C mismatches occurred, based on the Poisson model. Clusters of reads corresponding to peaks were then annotated using the ENSEMBL v112 gene transfer format (GTF) annotation (24). After determining the proportions of read clusters mapping to protein-coding and non-coding regions, only DDX3X-crosslinked reads originating from clusters with valid annotations mapping to exons of protein-coding genes were considered for further analysis.

For RNAseq, reads were aligned to the genome using STAR (v 2.7.10b) (21), and gene counts were generated from the resulting binary alignment map (BAM) files using featureCounts (v2.0.8) (25).

For alignments to the SeV genome, a Cantell-strain genome FASTA sequence was used (ATCC). Viral transcription and translation start sites were identified using a custom GTF annotation based on an existing SeV annotation (NCBI RefSeq assembly GCF_000855625.1) with viral 5’ untranslated regions (UTRs) explicitly annotated. Human alignments were performed using the UCSC hg38 RefGene assembly (26).

### NGS Data Analysis

Transcript-specific 5’UTRs, start codons, and coding sequences were identified based on the UCSC RefGene annotation (26). Transcripts per million (TPM) values of transcript variants in the RNAseq data were quantified using Salmon (v1.10.1) (27) and averaged between all conditions. For each gene, the most abundant transcript variant was selected, and the spliced sequences of the 5’UTR and whole transcript were extracted using gffread (28). 5’UTR+ sequences were generated by truncating the transcript to the 5’UTR + start codon and 50 additional nucleotides of the CDS.

Metagene analyses were performed as in (29). Briefly, genomic positions of gene 5’UTRs, CDS and 3’UTRs were length-normalised to 11, 70, and 100 equally-sized bins, respectively. Clusters of DDX3X-crosslinked PAR-CLIP reads identified as binding sites from peak calling were then assigned to these bins based on their genomic coordinates, and normalised counts of clusters in these bins were plotted.

For analyzing 5-mers, protein-coding genes were designated as DDX3X-bound if the gene had at least one DDX3X PAR-CLIP T-to-C cluster mapping to an exon; otherwise, they were designated as unbound. The occurrence of each 5-mer within all 5’UTR+ sequences belonging to genes in either group was then counted and normalized to the total number of nucleotides in either group.

Predicted minimum free energy secondary structures and ΔG values were computed using the RNAfold tool of the Vienna RNAfold suite (30) with default parameters, using FASTA sequences of 5’UTR+ regions as input.

Figures of RNAseq and PAR-CLIP reads aligned to genes were generated using the Integrative Genomics Viewer (v2.13.1) (31). Protein interaction network and Gene Ontology (GO) analyses were performed using STRING (v12.0) (32).

### Reporter Construct Cloning and Luciferase Assays

5’UTR regions of interest were amplified from either gDNA or cDNA using Phusion high-fidelity polymerase (NEB, M0530S) according to the manufacturer’s instructions, using PCR primers with 15 nt homologous regions to the NcoI and HindIII restriction sites of the pGL3-Prom vector (Promega, E1761). Primer sequences are detailed in Table S2. Inserts were cloned into the vector using the In-Fusion® Snap Assembly kit (Takara Bio, 638943) and transformed into Stellar competent *E. coli* cells (Takara Bio, 636766) as per manufacturer’s instructions. All constructs were sequence-verified using Sanger sequencing (MWG Eurofins), and the sequences of inserts are detailed in Table S3.

For luciferase assays, NSC and shDDX3X HEK293T cells were seeded in a 96-well plate at a density of 10^5^ cells/well in full DMEM containing 2 µg/ml doxycycline to induce knockdown. The next day, cells were transfected simultaneously with equal amounts of pGL3-5’UTR-Firefly reporter construct and pGL3-Renilla (Promega, E2231) control, in addition to empty vector (pCMV-myc) for a total of 230 ng DNA per well, using 5% Lipofectamine (Invitrogen, 11668027) in serum-free DMEM. 64 hours post-seeding, wells were lysed with 50 µl reporter lysis buffer (Promega) at room temperature for 15 min, followed by freeze-thawing at -20°C. 20 µl of lysate was transferred to two separate opaque white flat-bottomed plates, to which 40 µl of either 0.1 µg/mL coelenterazine in PBS or luciferin substrate solution (100 mM Tricine, 12.5 mM MgSO_4_, 10 mM EDTA, 0.15 mM DTT, 2.5 mM ATP, 1.25 mM Acetyl-CoA, 2.25 mM Firefly Luciferin, 25 mM NaOH, 1.25 mM Magnesium Carbonate Hydroxide) was added, followed by immediate reading of luminescence at 480 nm or 580 nm for RLUC or FLUC activity, respectively. FLUC/RLUC ratios were taken for each well, the median ratio of a technical triplicate was taken, and, for each condition, normalized to a corresponding control sample transfected with pGL3-Prom with no insert.

### Statistical Analyses and Figures

Statistical analyses and figures were generated with R (v4.1.1) (33), using the DESeq2 (v1.34.0) (34), pheatmap (v1.0.12) (35) and ggplot2 (v3.5.0) (36) packages. For luciferase reporter assays, statistical significance was assessed using a two-sided Wilcoxon rank-sum test paired by biological replicate. For RNAseq, differential gene expression analysis was performed using DESeq2 (v1.34.0) (34), and statistical significance was assessed using the default Wald Test with p values corrected using the Benjamini-Hochberg method. In all analyses, a p-value < 0.05 was considered statistically significant. Code for the generation of most figures related to NGS data can be found on GitHub (https://github.com/cathalsr/ddx3x_sev).

### Western Blotting

Protein samples were subjected to SDS-PAGE using 10% bis-tris polyacrylamide gels. Proteins were transferred to PVDF membranes using a BioRad Trans Blot Turbo with constant 1.25 A/membrane for 30min with gels and membranes soaked in transfer buffer (48 mM Tris, 39 mM glycine, 20% methanol). Membranes were then incubated in blocking buffer (5% skimmed milk powder (Marvel), 0.1% Tween-20 in PBS) for 1 h, then incubated overnight with 3mL of antibody solution (0.1µg/mL of anti-DDX3X mAb (Santa Cruz Biotechnology, 2253C5A), 1µL/mL anti-SeV pAb (MBL, PD029), or 0.1µg/mL anti-beta actin (Sigma Aldrich, A5441) in blocking buffer) at 4°C. Membranes were washed three times for 5min each in wash buffer (0.1% Tween-20 in PBS) before incubation with 3 mL secondary antibody solution (1µg/mL anti-mouse-HRP (Thermo Fisher Scientific A16084), or 1 µg/mL anti-rabbit-HRP (Sigma Aldrich, A8275) in blocking buffer) for 2 h at RT. Membranes were then again washed three times for 10 min each. ECL substrate (2.5mM luminol, 0.396 mM p-coumaric acid, 100 mM Tris-HCl pH 8.5, 0.000183% H_2_O_2_) was added to the surface of the PVDF, and blots were imaged using a Syngene G:Box imager (Syngene) or an iBright CL1500 Imaging system (Invitrogen). Band quantification was performed using iBright Analysis Software, with per-sample intensities normalized to corresponding b-actin band intensities.

### ELISA

For ELISAs, undiluted cell culture supernatants collected from NSC and shDDX3X HEK293T cells after 72 h knockdown and (where indicated) 16hpi with SeV were assayed using a Human IFN-beta ELISA kit (R&D systems, 41410-1), following manufacturer’s instructions.

## RESULTS

### DDX3X binds structured mRNAs during SeV infection

Despite the strong links between DDX3X and many different viruses (7), it is currently unknown whether and how viral infections affect DDX3X’s function as a regulator of host mRNA translation. We therefore set out to investigate whether infection with Cantell-strain Sendai Virus (SeV), a model RNA virus that strongly activates the RIG-I pathway and induces robust IFNB expression (37), alters DDX3X’s interactions with host RNAs either directly or via the resulting anti-viral host response. To this end, we employed the PAR-CLIP method (18) on HEK293T cells which were either left uninfected or infected for 2 or 16 hours with SeV to identify DDX3X mRNA binding sites throughout the entire transcriptome at nucleotide resolution.

Immunoprecipitation of endogenous DDX3X from 4-thiouridine-treated and UV-irradiated cells achieved efficient and specific pulldown of DDX3X protein (Fig. S1A). Detection of fluorescence from SDS-PAGE gels revealed a band corresponding to crosslinked DDX3X-RNA complexes with an apparent molecular weight approximately 25kDa greater than DDX3X protein (Fig. S1A). This was specific to pulldown with anti-DDX3X antibody, as this band was not detected when an anti-HNRNPK antibody was used for immunoprecipitation (Fig. S1A). This region was excised and further processed for PAR-CLIP library generation and sequencing. Detection of T-to-C conversions at cross-linking sites induced by PAR-CLIP allowed for identification of high-confidence binding sites via peak calling, corresponding to ∼30,000-100,000 binding sites mapping to ∼8,000-14,000 unique annotated genes per-sample (n=3 for uninfected and 2hpi, n = 2 for 16hpi) (Fig. S1B, S1C). For all samples, DDX3X binding occurred predominantly within exons of protein-coding mRNAs (Fig. S1D). A notable minority of reads mapped to intronic regions, perhaps as a result of alternative splicing leading to retained introns within DDX3X-bound mRNAs, or possibly due to a minority population of nuclear DDX3X binding pre-mRNA being detected. Among mRNAs identified as DDX3X bound, the majority were detected in both infected and uninfected samples, with a higher number of mRNAs uniquely identified in uninfected samples (Fig. 1A), although this is likely due to the higher overall number of DDX3X-bound transcripts identified in these samples (Fig. S1B).

**Figure 1:**
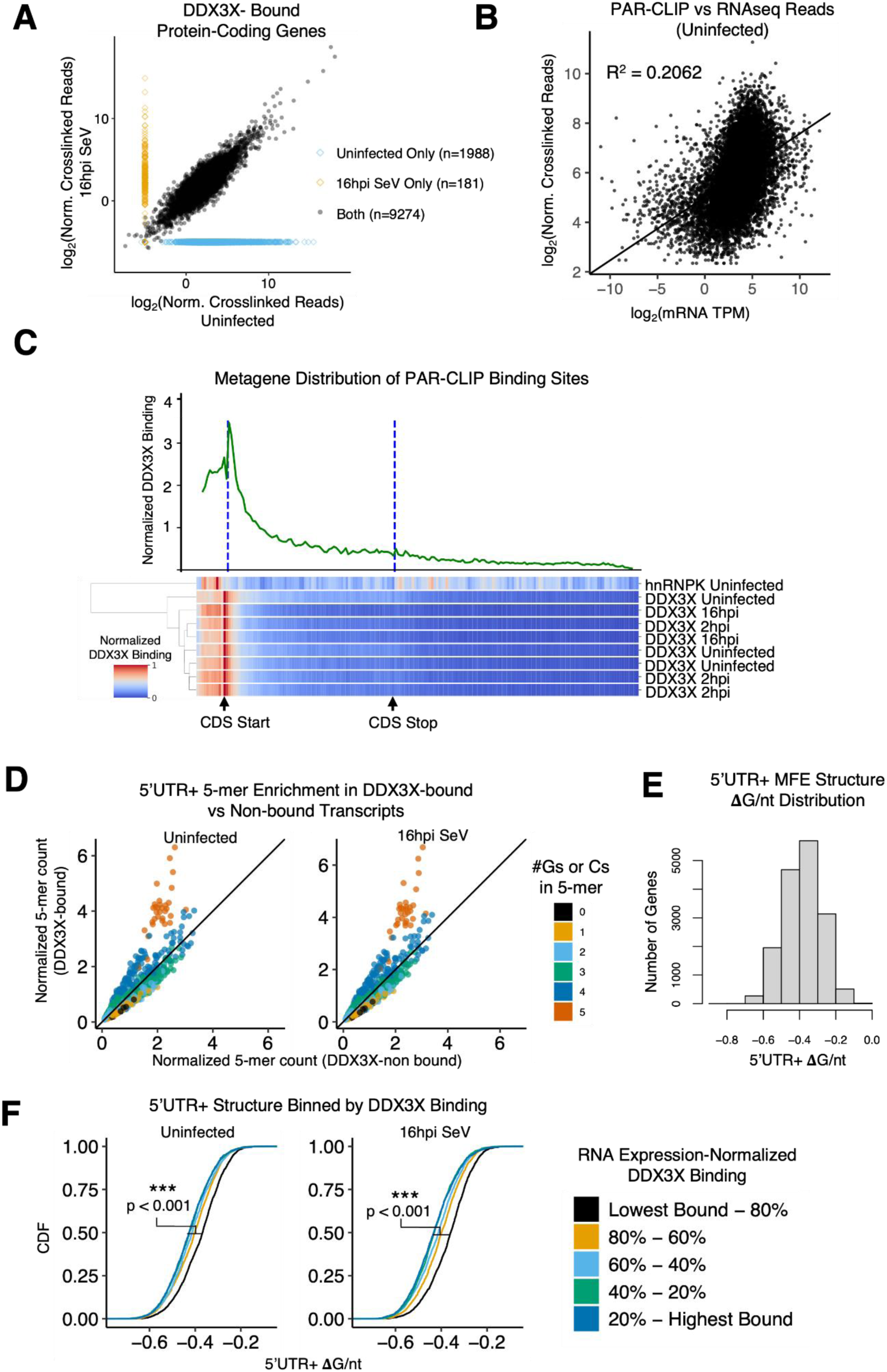
DDX3X preferentially binds structured 5’UTRs and upstream CDS regions irrespective of SeV infection. (A) Plot of per-gene mean normalized DDX3X-crosslinked read counts (log_2_(#reads per-gene / million total reads per-sample)) in uninfected vs. 16 hours post infection (hpi) SeV samples. (B) Plot of per-gene mean DDX3X-crosslinked reads (normalized to total number of DDX3X-crosslinked reads per-sample) from binding sites in exons of protein-coding genes from PAR-CLIP versus per-gene average transcripts per million (TPM) values from RNAseq. (C) (Top) Example metagene plot of a representative DDX3X PAR-CLIP sample (uninfected), where crosslinked reads from T-to-C clusters of protein-coding transcripts were assigned to bins mapping to 5’UTR, CDS and 3’UTR regions. (Bottom) Hierarchically-clustered metagene heatmap comparing DDX3X (SeV infected and uninfected) and HNRNPK PAR-CLIP data. (D) Enrichment of 5-mers of different GC content between DDX3X bound and non-bound 5’UTR+ regions. Per-gene 5’UTR+ sequences of the most abundant transcript variants detected in HEK293Ts via RNAseq were assigned as either DDX3X-bound or non-bound based on the presence or absence of DDX3X PAR-CLIP T-to-C clusters. The abundance of each 5-mer was then counted in both groups, and normalized to the total nucleotide content in both groups. (E) Histogram of ΔG/nt values of *in silico* predicted minimum free energy (MFE) structures of 5’UTR+ regions for all genes expressed in HEK293T cells based on our RNAseq data (F) Cumulative distribution function (CDF) plots of predicted ΔG/nt values, binned by level of DDX3X binding in uninfected and 16h SeV samples. P values are from a Mann-Whitney U test comparing the distributions of ΔG/nt values in the highest-bound vs lowest-bound bin. * = p < 0.05, ** = p < 0.01, *** = p < 0.001.

Based on this data and previously published CLIP studies (3, 9, 38, 39), DDX3X associates with a large proportion of the transcriptome, and it has so far proven difficult to identify specific determinants of DDX3X binding, such as RNA sequence motifs (3). Thus, the question could arise whether DDX3X promiscuously binds to all mRNAs, where mRNA expression level is the main determinant of DDX3X binding level. To address this question, we plotted the number of DDX3X crosslinked reads belonging to high-confidence binding sites against mRNA levels for each gene, which showed poor correlation (Fig. 1B), suggesting that mRNA abundance alone is not a strong predictor of DDX3X binding.

To further examine DDX3X’s binding distribution on its target mRNAs, we next performed a metagene analysis of DDX3X-crosslinked reads aligning to protein-coding genes. The 5’UTR, CDS, and 3’UTR coordinates of genes were length-normalized to a number of equally-sized bins to which DDX3X binding sites were assigned. This showed that binding of DDX3X to transcripts was predominantly enriched within the 5’UTR, peaking proximal to the start codon and extending noticeably into the adjacent early CDS (Fig. 1C), a pattern broadly in agreement with previously published DDX3X CLIP-seq studies (3, 9, 38, 39). For comparison, we included PAR-CLIP data of the unrelated RBP hnRNPK in this analysis, which did not show a similar enrichment around the start codon (Fig. 1C). There was no apparent difference in the metagene profiles generated from uninfected and infected HEK293T samples, and hierarchical clustering of metagene distributions of the different PAR-CLIP samples did not substantially segregate infected and uninfected samples (Fig. 1C), suggesting that SeV infection does not dramatically alter DDX3X’s distribution on target mRNAs.

Although DDX3X has so far mainly been suggested to regulate mRNA translation through binding to 5’UTRs, the binding peak we observed stretched substantially into the early CDS (Fig. 1C) and we reasoned that this region may also influence translation initiation regulation due to proximity to the start codon. Therefore, we defined a region encompassing the 5’UTR, start codon, and the first 50 nt of the CDS (hereafter named “5’UTR+ region”) to be used in further analyses.

DDX3X has previously been shown to preferentially bind GC-rich, highly structured sequences (3, 38). To determine if this feature of DDX3X binding is reproducible in our data, we quantified the occurrence of all possible 5-mers in 5’UTR+ regions within transcripts expressed in HEK293T cells and compared between DDX3X-bound and non-bound transcripts. Compared to non-bound transcripts, the 5’UTR+ regions of DDX3X-bound transcripts were highly enriched for 5-mers containing predominantly Gs and Cs, in broad agreement with previous studies (Fig. 1D). This enrichment was not altered after SeV infection (Fig. 1D), suggesting that DDX3X preference for GC-rich 5’UTR+ regions is not affected by infection.

High GC-content tends to be associated with the occurrence of more stable secondary structures in RNAs (40), and resolution of mRNA secondary structures has been highlighted as the main mechanism underlying DDX3X-mediated translation regulation (7, 11). Thus, to more directly examine the relationship between DDX3X binding and RNA secondary structure, we first generated the minimum free energy structures (MFE) and corresponding delta free energy (ΔG) values of 5’UTR+ regions for HEK293T-expressed genes *in silico* using Vienna RNAfold (30), with lower ΔG values indicating more thermodynamically stable secondary structures. The resulting length-normalized ΔG (ΔG/nt) for 5’UTR+ sequences were roughly symmetrically distributed with a median ΔG/nt of -0.39 (Fig. 1E). When 5’UTR+ ΔG values were binned by their respective gene’s number of mRNA level-normalised DDX3X-crosslinked reads, the degree of 5’UTR+ secondary structure increased progressively with the level of DDX3X binding (Fig. 1F). This data demonstrates that DDX3X binding was significantly enriched on transcripts with more structured 5’UTR+ regions compared to poorly-structured transcripts. The fact that we observed this in both infected and uninfected cells (Fig. 1F) again demonstrates an unaltered binding preference of DDX3X in virus-infected compared to uninfected cells.

In summary, we robustly determined DDX3X binding sites across the transcriptome in both uninfected and SeV-infected HEK293T cells (Fig. 1A, S1B, S1C). We found that DDX3X preferentially binds transcripts featuring GC-rich 5’UTR+ regions displaying a high degree of secondary structure in line with previous studies, and demonstrated that these binding characteristics are not significantly altered by SeV infection.

### SeV infection changes the DDX3X target pool

Although it displayed the same general binding characteristics in SeV-infected cells compared to uninfected cells, we reasoned that DDX3X’s interactions with the host transcriptome are likely still altered by infection, even simply through host transcriptome changes in response to infection. To determine SeV-induced transcriptome changes, RNAseq data of HEK293T cells at 2h and 16h post SeV infection were compared to those of uninfected cells. SeV infection at 16 hpi resulted in robust induction of *IFNB* and Interferon-Stimulated Genes (ISGs) as expected, while at 2 hpi no significant transcriptome changes were observed (Fig. 2A), corresponding to previously observed kinetics for the RIG-I/type I IFN response to SeV (41). To identify mRNAs that are upregulated in response to SeV infection and also bound by DDX3X, changes in the number of DDX3X PAR-CLIP crosslinked reads per gene were plotted against their corresponding mRNA level changes measured via RNAseq. As shown in Figure 2B, many but not all mRNAs that were strongly induced 16 hpi, including mRNAs for several ISGs and *IFNB* itself, were bound by DDX3X. To further examine these putative DDX3X targets, the STRING database (32) was used to perform a Cluster and Gene Ontology (GO) analysis on the subset of genes that was both upregulated at the mRNA level after infection and detected as DDX3X-bound exclusively in infected conditions. This revealed a network of genes (Fig. 2C) with strong enrichment for GO terms relating to viral infection and the anti-viral innate immune response (Fig. 2D), highlighting these pathways as novel areas of DDX3X post-transcriptional regulation. Because these genes are not or only very lowly expressed in uninfected cells, previous CLIP-Seq studies on DDX3X conducted in uninfected cells could not have detected these interactions.

**Figure 2:**
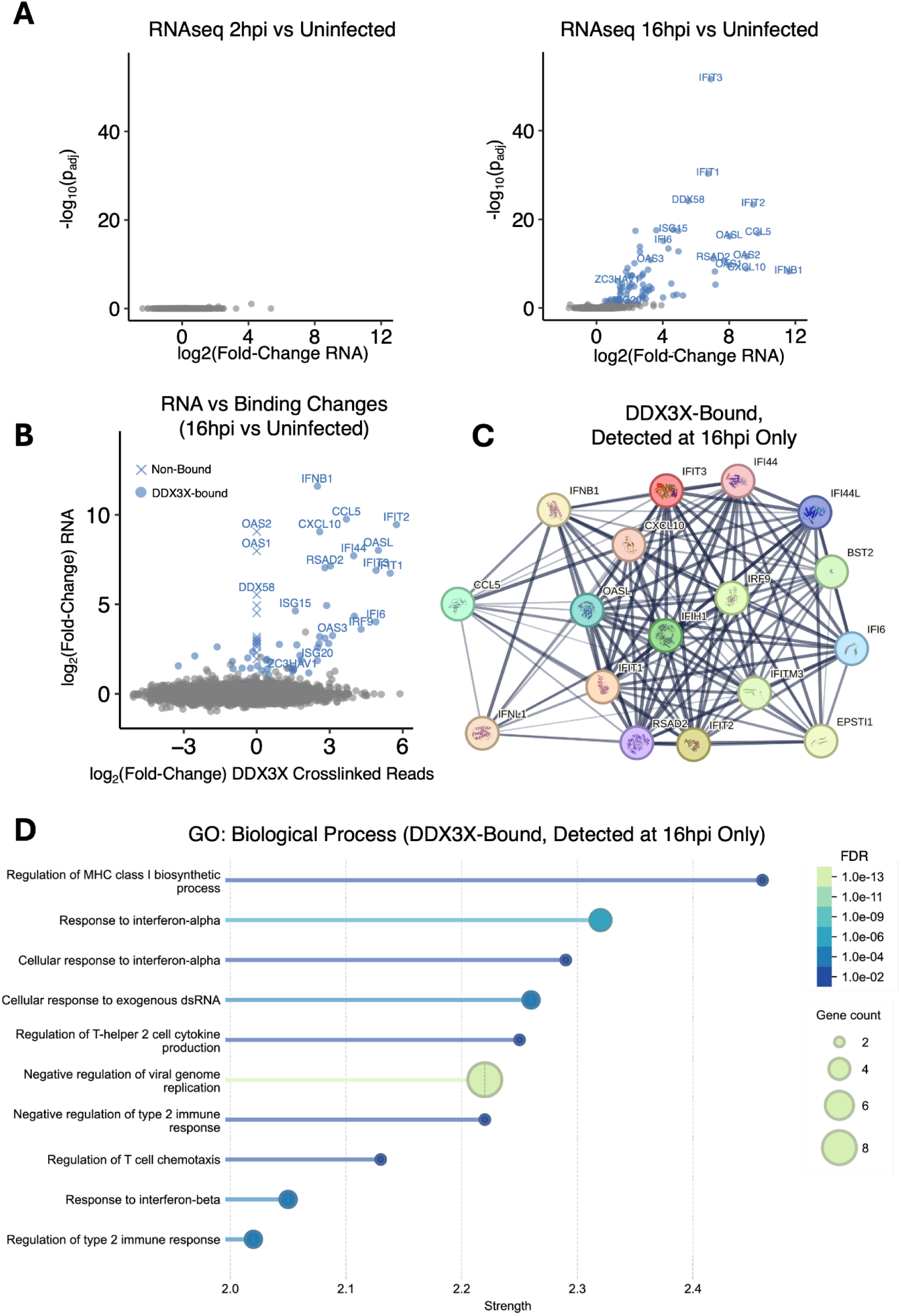
SeV infection alters the DDX3X mRNA target pool. (A) Volcano plots of RNA expression level changes in 2hr and 16h SeV-infected vs uninfected cells, determined via RNAseq (uninfected & 16hpi infected n = 6, 2hr infected n = 4). Genes highlighted in blue have a padj < 0.05, and the names of IFNB and some ISGs are shown. (B) Per-gene log2-fold change in DDX3X-crosslinked reads versus log2-fold change in mRNA comparing 16hpi vs uninfected cells. Genes highlighted in blue have significantly altered mRNA expression levels (padj < 0.05), and the names of IFNB and some ISGs are shown. Non-DDX3X bound transcripts are shown with a fold change in DDX3X-crosslinked reads of 0. (C) Protein interaction network for genes with significant mRNA level changes at 16hpi that were detected as DDX3X-bound exclusively at 16hpi, generated by STRING (32). (D) Enrichment of Gene Ontology: Biological Process terms for genes in (C) as provided by STRING (32). Strength = log_10_(#observed genes per-term / #expected genes per-term with a randomized sample).

### *IFNB* mRNA translation is regulated by DDX3X

We were particularly interested in the novel finding that DDX3X binds to the *IFNB* mRNA because of its key function as an anti-viral cytokine and our previous work showing that DDX3X acts as a signaling adaptor in the RIG-I/IRF3 pathway that upregulates *IFNB* transcription (16, 17). DDX3X’s binding to the *IFNB* mRNA (Fig. 2B, 2C) suggests that it may additionally regulate *IFNB* expression post-transcriptionally. We therefore closely examined the distribution of DDX3X-crosslinked reads mapping to the *IFNB* mRNA in samples from cells harvested at 16 hpi. Although crosslinked reads were mostly proximal to the start codon in a manner similar to the metagene distribution (Fig. 1C), the actual start codon itself showed poor coverage (Fig 3A). Instead, we observed a roughly bimodal pattern of two binding peaks flanking the start codon in both the 5’UTR and the start of the CDS. Based on the MFE structure predicted for the *IFNB* mRNA using Vienna RNAfold (30), the two DDX3X binding sites appear to correspond to the base of a stem loop containing the start codon (Fig. 3B). We reasoned that this stem loop could conceivably exert an effect on IFNB translation initiation that is modulated by DDX3X.

**Figure 3:**
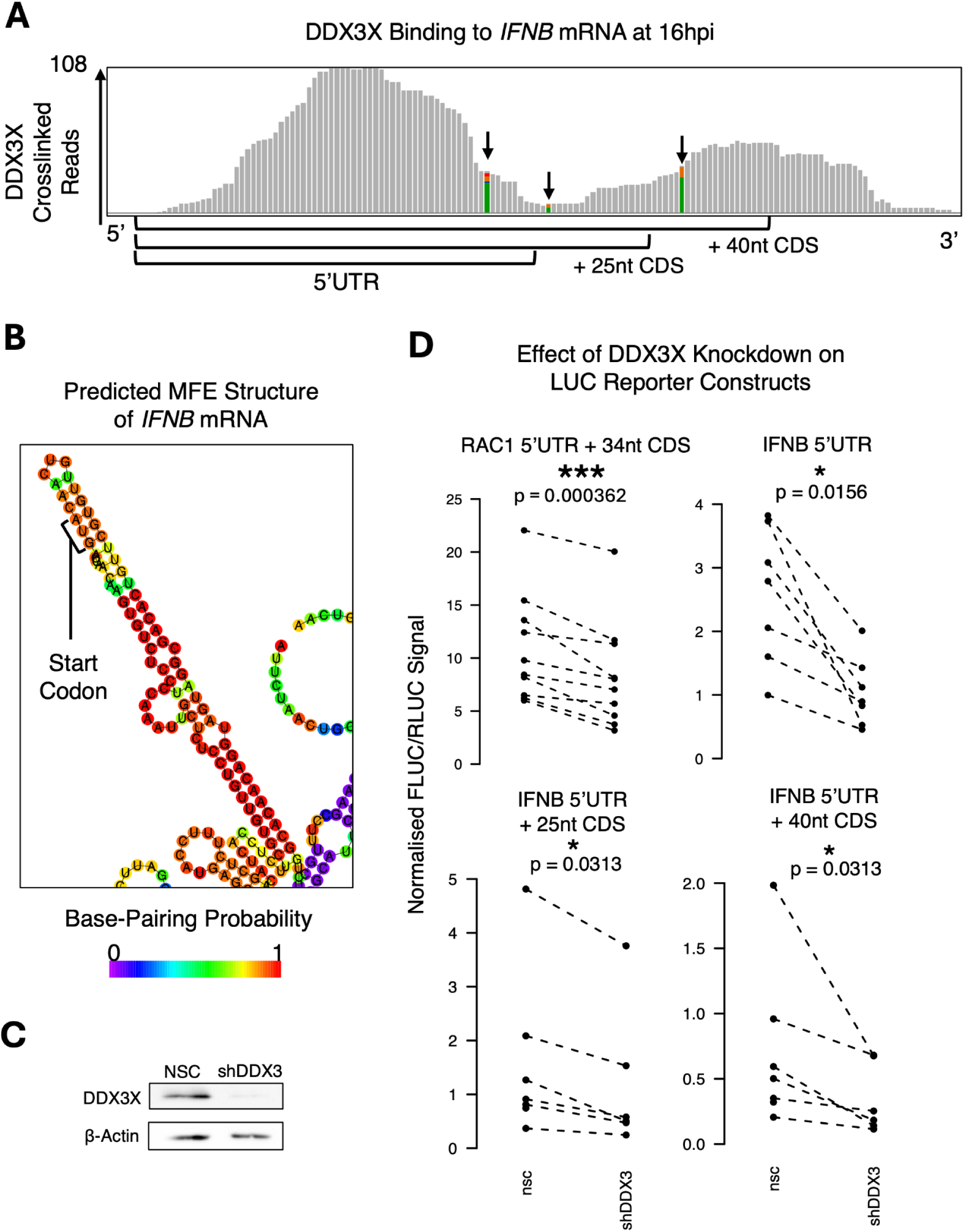
DDX3X binds and regulates translation of IFNB mRNA. (A) Interactive Genomics Viewer display of the distribution of DDX3X-crosslinked reads proximal to IFNB1 start codon in PAR-CLIP data from HEK293T cells 16hpi (one sample shown of n = 2). Sites of PAR-CLIP-induced T to C conversions (represented as A to G mismatches in IGV for an antisense transcript) are highlighted, where the ratio of green:orange indicates the ratio of A:G among reads aligning to a nucleotide position. (B) Detail from the predicted minimum free energy (MFE) structure of IFNB1 mRNA as predicted by Vienna RNAfold (Fig. S2A), displaying a local secondary structure enclosing the start codon, and occurring proximal to the predicted DDX3X binding site. (C) Western blot of HEK293T NSC and shDDX3X cell lysates after 72h of doxycycline treatment, confirming successful knockdown induction. (D) Effects of DDX3X knockdown on FLUC/RLUC signal from cells transfected with pGL3-Prom Firefly luciferase 5’UTR reporter constructs and pGL3-Renilla luciferase. The FLUC signal was normalised to RLUC signal for each sample, and median values of triplicates were then normalised to a pGL3-Prom control with no 5’UTR insert. Reporters contained either RAC1 5’UTR and 34nt of the CDS, or IFNB 5’UTR with varying lengths of the CDS, and were transfected into non-silencing control (NSC) or short-hairpin DDX3X knockdown (shDDX3X) HEK293T cells treated with doxycycline for 72h to induce DDX3X knockdown. P values are calculated via Mann-Whitney U test paired by biological replicate. * = p < 0.05. *** = p < 0.001.

To determine whether DDX3X binding to this region is relevant for IFNB translation regulation, the *IFNB* 5’UTR combined with varying lengths of the adjacent CDS (0, 25, 40nt), was PCR-amplified and inserted upstream of the firefly luciferase (FLUC) coding sequence into the pGL3-Prom vector to generate 5’UTR reporter gene constructs (Fig. 3B). Of the different variations of the *IFNB* reporter, the longest, containing the start codon plus 40 nt of the CDS, is predicted to reproduce the DDX3X-bound stem loop which appears in the predicted MFE structure of the whole *IFNB* mRNA (Fig. 3B, S2). To test the effects of DDX3X manipulation, we transfected these reporters into HEK293T cell lines stably expressing inducible DDX3X knockdown (shDDX3X) or a non-silencing control (NSC) shRNA that were previously described (15), and co-transfected a pGL3-Renilla luciferase (RLUC) construct to normalize reporter FLUC signal to RLUC signal for each sample. As a positive control, we also tested a reporter containing the 5’UTR and early CDS of *RAC1*, an mRNA well-known to be subject to DDX3X-mediated translation regulation (3, 6).

In the reporter assay samples, treatment of cells with doxycycline resulted in knockdown of DDX3X in shDDX3X compared to NSC cells, as confirmed by western blot (Fig. 3C), and the knockdown resulted in decreased signal of the RAC1 reporter, indicating decreased translation as expected (Fig. 3D). The *IFNB* 5’UTR with 40nt of the CDS also displayed DDX3X sensitivity (Fig. 3D), suggesting that the DDX3X interaction with the 5’UTR-CDS spanning stem loop regulates translation of *IFNB*. However, we also observed similar DDX3X sensitivity for reporters containing the *IFNB* 5’UTR in isolation or with only 25 nt of the CDS (Fig. 3D), which may suggest that the 5’UTR alone is sufficient for DDX3X-mediated translation regulation of *IFNB*.

In summary, we observed DDX3X binding to both the 5’UTR and early CDS of *IFNB*, coinciding with the presence of a predicted stem loop enclosing the IFNB translation start codon. We also demonstrated that DDX3X depletion decreases translation of reporters containing the *IFNB* 5’UTR, suggesting that the *IFNB* mRNA is indeed regulated by DDX3X at the level of translation initiation.

### DDX3X interacts with SeV mRNA

Although DDX3X has been shown to interact with RNA species from a diverse range of viruses, (7) and SeV infection has been used previously to identify DDX3X’s function in the RIG-I pathway (17), it is currently unknown whether DDX3X directly interacts with SeV and regulates its replication. We therefore next examined whether DDX3X binds to SeV RNA by aligning DDX3X PAR-CLIP reads to the Cantell strain SeV genome.

As a negative-stranded RNA virus, SeV produces various RNA species over the course of its replication cycle, mainly negative-sense genomic RNA, positive-sense antigenomic RNA, and positive-sense viral mRNA transcripts (Fig. 4A). PAR-CLIP is capable of identifying whether RBP-crosslinked reads are from positive or negative-strand transcripts, so the type of SeV RNAs bound by DDX3X may be at least partially inferred from this information. Therefore, we first attempted to discern which type of viral RNAs DDX3X bound in infected conditions.

**Figure 4:**
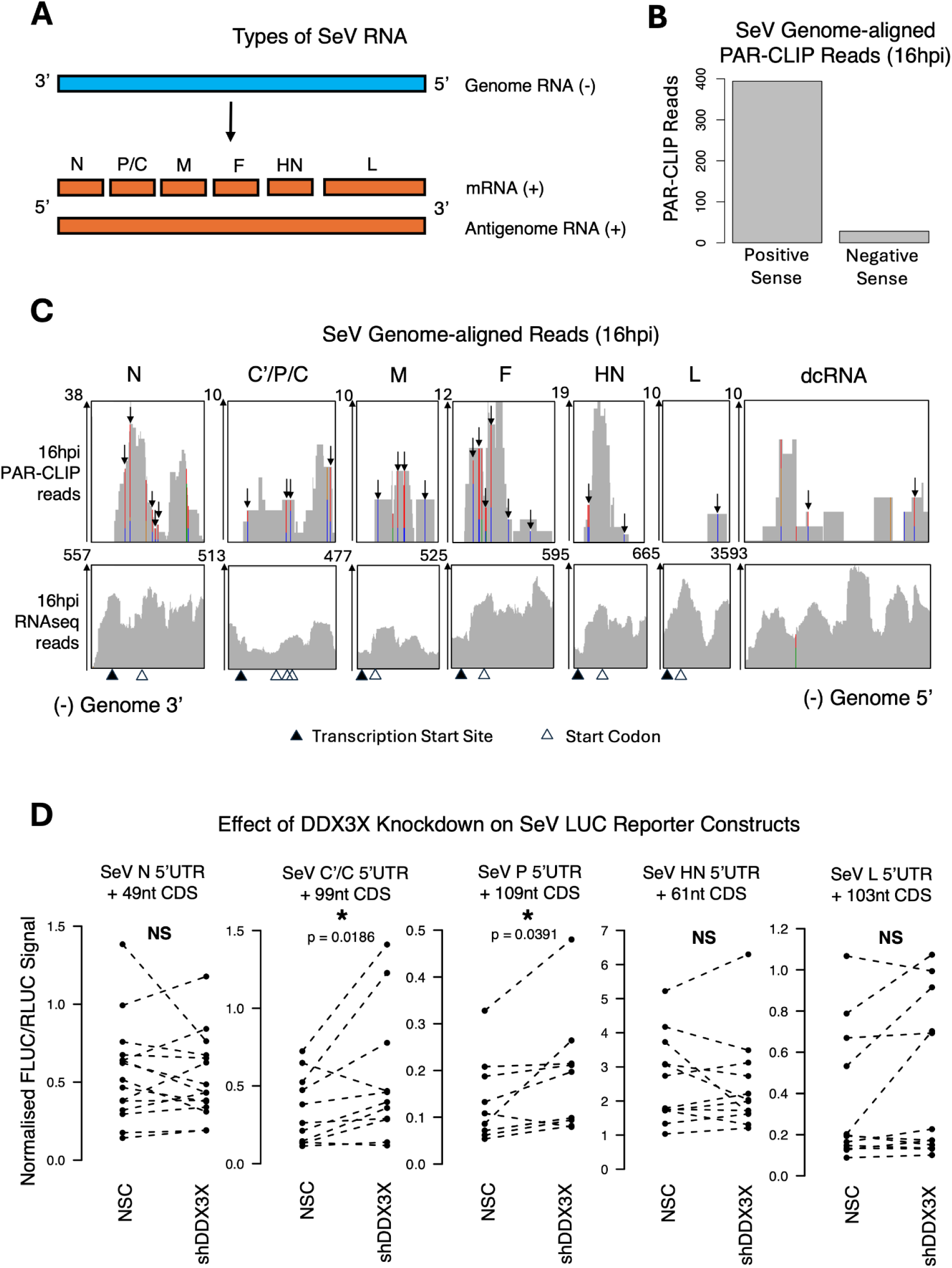
DDX3X interacts with positive-strand SeV RNA. (A) Overview of the different SeV RNA species. (B) Mean count of forward vs. reverse reads in 16hr SeV infected DDX3X PAR-CLIP samples aligned to SeV Cantell strain genome. (C) Distribution of RNAseq reads and DDX3X-crosslinked PAR-CLIP reads aligning to regions of the SeV Cantell strain genome at 16hpi (single RNAseq replicate of n = 6 and PAR-CLIP replicate of n = 2 shown). Transcription start sites and start codons of SeV genes, as well as a region presumed to correspond to SeV dcRNA, are shown. PAR-CLIP T-to-C conversions are indicated as T-to-C mis-matches. Colored sites possess a large proportion of reads with nucleotide mismatches, with the ratio of A,T,C,G shown by the ratio of green, red, blue and orange, respectively. (D) Effects of DDX3X knockdown on FLUC/RLUC signal from cells transfected with pGL3-Prom Firefly luciferase 5’UTR reporter constructs and pGL3-Renilla luciferase. The FLUC signal was normalised to RLUC signal for each sample, and median values of triplicates were then normalised to a pGL3-Prom control with no 5’UTR insert. Reporters containing SeV 5’UTRs were transfected into non-silencing control (NSC) or short-hairpin DDX3X knockdown (shDDX3X) HEK293T cells treated with doxycycline for 72h to induce DDX3X knockdown. P values are calculated via Mann-Whitney U test paired by biological replicate. * = p < 0.05, NS = p > 0.05.

Aligned reads detected at 16 hpi were predominantly positive-sense (Fig. 4B), suggesting that DDX3X binds to SeV antigenomic and/or mRNA rather than genomic RNA. Of note, the number of DDX3X-crosslinked reads in the PAR-CLIP data was low compared to most host transcripts, despite an abundance of SeV RNA reads in cells at 16 hpi detected in our RNAseq data (Fig. 4C). This suggests that DDX3X is not strongly re-directed to SeV RNAs during infection, and thus viral RNAs are unlikely to compete with host transcripts for DDX3X binding.

In the RNAseq data, we also observed a region at the genome 5’ end displaying a drastically greater abundance of aligned reads than the rest of the genome (Fig. 4C), which likely corresponds to defective copyback genomic RNA (dcRNA), an immunostimulatory dsRNA produced as a byproduct of SeV replication known to be abundant in Cantell strain SeV (Sánchez-Aparicio et al., 2017). Despite the abundance of RNAseq reads aligning to the dcRNA region compared with the rest of the SeV genome, very few PAR-CLIP reads were detected in this region (Fig. 4C), suggesting that DDX3X does not readily bind SeV dcRNAs.

Interestingly, the majority of PAR-CLIP reads with T-to-C conversions occurred in proximity to viral start codons, in a manner strikingly similar to DDX3X’s binding pattern with host RNAs (Fig. 1C). Such binding, albeit weak, was observed for all SeV genes with the exception of SeV L (Fig. 4C).

To examine potential DDX3X-mediated translation regulation of SeV genes, pGL3-Prom 5’UTR luciferase reporter gene constructs for SeV N, HN, L, C’/C and P were created. Similar to how we designed the IFNB reporter gene constructs, we included parts of the early CDS with the 5’UTR into our SeV reporters to capture regions of DDX3X binding downstream of the translation start codon (Fig. 4C). Since it displayed negligible binding by DDX3X (Fig. 4C), L was included as a negative control. The C/P gene is notable in having multiple translation initiation sites (42). Thus, two reporter constructs were made for this gene, one with the luciferase CDS in-frame with the P initiation site, and the other in-frame with the C’ and C initiation sites (C’/C). The SeV 5’UTR reporter gene constructs were then tested in HEK293T cell lines expressing either a DDX3X shRNA or a non-silencing control (NSC) shRNA as described above for IFNB. Upon DDX3X knockdown, the reporters showing the most consistent change in activity were C’/C and P, which displayed increased reporter activity (Fig. 4D), suggesting that DDX3X may negatively regulate translation of both C’/C and P. Despite showing the most distinct DDX3X binding peaks around the translation start codon among the SeV genes, we did not detect significant differences in reporter activity in response to DDX3X knockdown for the SeV N and HN genes (Fig. 4D).

In summary, DDX3X appears to bind several SeV mRNAs, but not strongly compared to host mRNAs. Its binding is therefore unlikely to be redirected from host to virus mRNAs during infection. Our 5’UTR reporter results suggest that DDX3X can negatively regulate translation from the C’/P/C 5’UTR, so it potentially has differential effects on host and viral transcripts.

### DDX3X knockdown results in decreased IFNB production but does not significantly impact SeV protein levels

The data from this study and our previous work (Schröder et al., 2008) points towards a consistent anti-viral role for DDX3X during SeV infection, via positive effects on both transcription and translation of the critical anti-viral cytokine IFNB and negative regulation of the SeV C’/P/C 5’UTR. Yet, DDX3X has been implicated as a positive host factor for many other viral infections, and we cannot rule out that DDX3X has additional effects on the viral life cycle or the cellular environment that affect the outcome of the viral infection. We therefore wanted to assess the overall effect of DDX3X depletion on the course of SeV infection.

To assess the total impact of DDX3X depletion on IFNB protein levels in response to infection, the levels of secreted IFNB from SeV-infected DDX3X shRNA and NSC HEK293 cells were assessed by ELISA. At 16 hpi, DDX3X knockdown cells displayed reduced secreted IFNB protein levels by roughly half (median IFNB concentration in knockdown relative to control = 0.52, Mann Whitney U test p = 0.016) (Fig. 5A), further confirming that DDX3X is required for efficient IFNB protein production during SeV infection.

**Figure 5:**
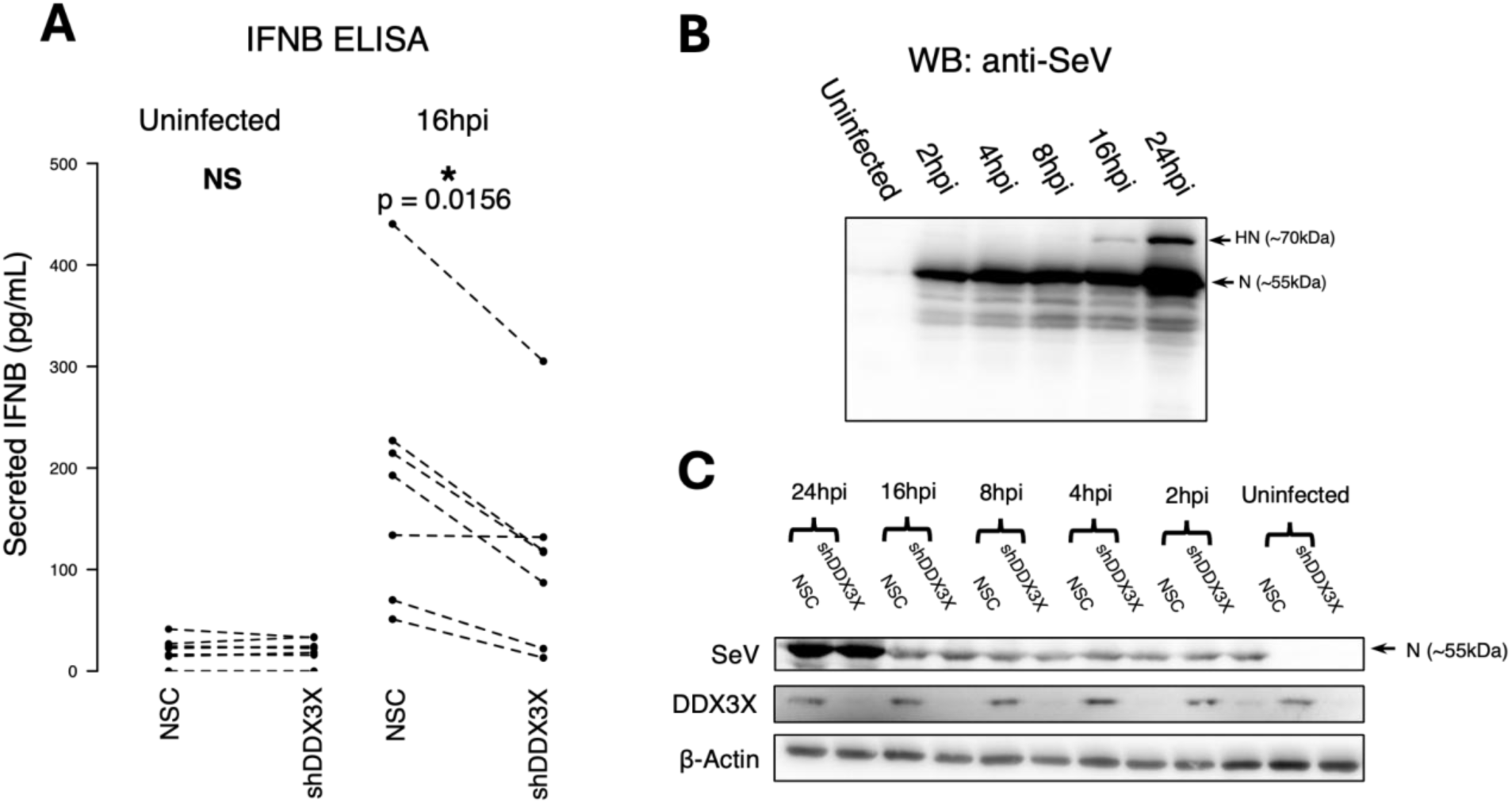
DDX3X knockdown leads to reduced IFNB protein production but does not affect SeV protein production during infection. (A) Levels of IFNB secreted by NSC and shDDX3X HEK293T cells either uninfected or infected for 16h with SeV, with 72h DDX3X knockdown, measured via ELISA (n = 6). DDX3X knockdown and even levels of β-actin between NSC and shDDX3X samples were confirmed via western blot of cell lysates as shown in (C). (B) Western blot of HEK293T cells infected with SeV for varying lengths of time, probed with anti-SeV polyclonal antibody. Approximate apparent molecular weights, as well as predicted protein identities of bands, are shown. (C) Western blot using anti-SeV pAb, DDX3X and β-actin antibody on samples from DDX3X knockdown and control HEK293Ts infected for various times with SeV. The sample shown in the representative blot corresponds to one of the biological replicates in (A). 2-3 biological replicates (dependent on time point) were carried out and quantified as shown in Figure S3.

One could expect that reduced IFNB production and/or enhanced C’/C/P viral protein production would lead to overall enhanced levels of SeV replication in infected HEK293T DDX3X shRNA compared to NSC cells. To examine this, we infected DDX3X shRNA and NSC HEK293T cells with SeV for varying durations and then examined SeV protein levels using a commercially available polyclonal antibody raised against SeV in Western Blots. As expected, SeV protein production increased with infection duration, suggesting active translation of viral mRNAs in infected cells (Fig. 5B). A band was detected at ∼70 kDa 16 h and 24 h post infection, which we assumed to correspond to SeV HN based on its apparent molecular weight (43). The strongest band we detected was at ∼55 kDa and likely corresponds to SeV N (Fig. 5B) (44). This SeV N band was consistently detected throughout the time course, present at the earliest timepoint (2h) and progressively increasing, with the strongest expression seen after 24h (Fig. 5B). We used quantification of this band, relative to β-actin, as a measure of overall SeV protein synthesis over the course of infection, as previous studies using this antibody have done (45, 46). However, we did not detect a consistent difference in SeV N protein levels between control and DDX3X knockdown cells at any of the observed time points (Fig. 5C, S3), suggesting that decreased IFNB production and increased translation of C’/C and P resulting from loss of DDX3X did not impact bulk viral protein synthesis under the examined conditions.

In summary, DDX3X knockdown significantly decreased secreted IFNB protein in response to SeV infection as expected (Fig. 3D), confirming its positive regulation of this critical anti-viral cytokine. However, this did not have an appreciable impact on SeV protein production in our experimental system. On the other hand this data also suggest that DDX3X does not have an additional pro-viral role during SeV infection that affects the accumulation of viral proteins by 24hpi.

## DISCUSSION

In this study, we comprehensively characterized DDX3X binding to cellular and viral mRNAs in SeV-infected cells and identified novel infection-induced DDX3X targets with roles in anti-viral immunity.

Transcriptome-wide, DDX3X binding was enriched on GC-rich, structured 5’UTRs and adjacent coding regions, consistent with previous studies (3, 9, 38, 39) and its role in regulating translation initiation. It is currently unclear whether this stems from specific binding to higher-order structural features in preferentially-crosslinked RNAs, such as G-quadruplexes (47–49), or is merely due to DDX3X remaining associated for longer with more structured RNAs during the process of scanning and unwinding. It was notable in our data that DDX3X binding accumulated around annotated translation start sites, stretching across into the early CDS (Fig. 1C). This may indicate a role for it in a late translation initiation event, such as start codon recognition and the switch from initiation to elongation, with DDX3X dissociating soon after elongation commences (Geissler et al., 2012). Our study demonstrated that DDX3X’s RNA binding preferences and pattern are not significantly altered by SeV or the virus-induced innate immune response (Fig. 1C, 1D, 1F), suggesting that the underlying mechanisms of DDX3X RNA binding preferences remained largely intact in these conditions.

The observation of DDX3X binding both upstream and downstream of the start codon (Fig. 1C) led us to consider the importance of 5’UTR/CDS-spanning structural elements for the regulation of translation initiation by DDX3X. However, while we did observe a relationship between DDX3X binding and 5’UTR+ secondary structure (Fig. 1F) and 5’UTR/CDS-spanning structures are present in many DDX3X-bound mRNAs (Fig. S4, S5), our data does not currently allow us to conclude that 5’UTR/CDS-spanning structural elements are required for DDX3X sensitivity. For the IFNB mRNA, DDX3X bound both sides of a predicted stem loop encompassing the translation start codon (Fig. 3A). Despite this, our reporter data indicated that the IFNB 5’UTR alone is sufficient to confer DDX3X sensitivity (Fig. 3D). One possible explanation for this may be that alternative structures form in our reporter mRNAs between the IFNB 5’UTR and the FLUC CDS that also confer DDX3X sensitivity (Fig. S2B). Further mechanistic work is therefore still required to better understand determinants of sensitivity to DDX3X regulation and the relevance of the observed binding pattern stretching across translation start sites.

Importantly, our study identified IFNB as an mRNA transcript regulated by DDX3X at the level of translation initiation. To our knowledge, post-transcriptional regulation of IFNB expression and in particular the function of its 5’UTR is largely unexplored, and thus our results not only provide novel insights on DDX3X but also on the expression regulation of this important anti-viral cytokine. Pharmacological inhibition of DDX3X is being explored as an antiviral strategy due to its role as an essential host factor for many different viruses (14). However, our findings highlight an important additional consideration: inhibiting DDX3X’s helicase activity may inadvertently suppress translation of IFNB and other antiviral mRNAs, potentially weakening innate immune responses depending on the specific virus and the cellular context. Nonsense and missense germline mutations in the *DDX3X* gene can cause the neurodevelopmental condition DDX3X syndrome and somatic *DDX3X* mutations have also been identified in many different cancer types (8, 9, 50). Considering our data, these mutations likely impair the ability of affected cells to produce IFNB, and dysregulation of IFNB translation may therefore also be relevant in these disease contexts.

DDX3X binding to SeV mRNAs was detectable but limited, suggesting that viral RNAs are unlikely to outcompete host RNAs for DDX3X binding (Fig. 4B, 4C). Interestingly, the binding we did observe occurred primarily near viral start codons, mirroring DDX3X’s distribution on host mRNAs and suggesting similar engagement during translation initiation (Fig. 1C, 4C). Our 5’UTR reporter assays indicated that DDX3X negatively regulates translation from the C′/C/P 5′UTR, while its effects on other viral 5′UTRs were minimal (Fig. 4D). This suppression of C′/C/P translation may be physiologically meaningful, as the C proteins were shown to suppress type I IFN induction and signaling (37, 41, 51), so this could further enhance DDX3X’s positive regulatory effect on IFNB production during SeV infection.

Finally, despite measuring significantly reduced IFNB protein production in DDX3X-depleted SeV-infected cells as expected (Fig. 5A), we did not observe increased SeV protein accumulation as one might expect when anti-viral responses are impaired (Fig. 5C, S3). A possible explanation for this may be that HEK293T cells are highly permissive to viral replication (52), and due to this viral replication and protein accumulation may have already progressed to such an extent at 16 hpi that the anti-viral innate immune response mediated by IFNB could not make significant impact. In less-permissive cell types the balance between the onset of the anti-viral response mediated by IFNB and viral replication/protein production may be different. One study also suggested that IFNB may not impact SeV infection greatly, possibly due to the virus expressing negative regulators of IFN signaling (53). Importantly however, the lack of observed effect of DDX3X knockdown on SeV protein levels also indicates that DDX3X is not needed as an essential host factor for SeV replication, in contrast to many other viruses (7).

In conclusion, we have shown that SeV infection reveals a cohort of novel DDX3X mRNA targets in the form of IFNB and ISGs, indicating that DDX3X’s function as an RNA remodeling enzyme is relevant to the anti-viral host response. Specifically, we have demonstrated DDX3X’s effect on the post-transcriptional regulation of IFNB, a cytokine with broad-spectrum anti-viral effects that is universally produced during anti-viral innate immune responses.

## ACKNOWLEDGEMENTS

We thank all members of the Schröder and Hafner labs who supported this work, contributed ideas, and helped trouble-shoot experiments. We also thank Dr Sanjay Gupta, Dr. Stefan Juranek, and Prof. Katrin Paeschke for facilitating the earlier attempts at DDX3X PAR-CLIP. We gratefully acknowledge Dr. Stefania Dell’Orso, Faiza Naz, and Shamima Islam (NIAMS Genomics Technology Section) for high-throughput sequencing. This research was supported by the Intramural Research Program of the National Institutes of Health (NIH). The contributions of the NIH authors were made as part of their official duties as NIH federal employees, are in compliance with agency policy requirements, and are considered works of the United States government. However, the findings and conclusions presented in this paper are those of the authors and do not necessarily reflect the views of the NIH or the U.S. Department of Health and Human Services.

## AUTHOR CONTRIBUTIONS STATEMENT

CR conducted the majority of experiments and data analysis shown in the paper. DGA and AHP were involved in conducting the PAR-CLIP experiments and subsequent data processing and analysis. ES conducted reporter gene assay and Western Blot replicate experiments. MH facilitated and oversaw PAR-CLIP experiments in his lab and contributed his expertise to data analysis and interpretation. MS conceived and directed the study. CR, MS and MH co-wrote the manuscript, which all authors then edited and approved.

## REFERENCES

1. de Bisschop, G., Ameur, M., Ulryck, N., Benattia, F., Ponchon, L., Sargueil, B. and Chamond, N. (2019) HIV-1 gRNA, a biological substrate, uncovers the potency of DDX3X biochemical activity. Biochimie, 164, 83–94.

2. Song, H. and Ji, X. (2019) The mechanism of RNA duplex recognition and unwinding by DEAD-box helicase DDX3X. Nat. Commun., 10, 3085.

3. Calviello, L., Venkataramanan, S., Rogowski, K.J., Wyler, E., Wilkins, K., Tejura, M., Thai, B., Krol, J., Filipowicz, W., Landthaler, M., et al. (2021) DDX3 depletion represses translation of mRNAs with complex 5′ UTRs. Nucleic Acids Res., 49, 5336–5350.

4. Soto-Rifo, R., Rubilar, P.S. and Ohlmann, T. (2013) The DEAD-box helicase DDX3 substitutes for the cap-binding protein eIF4E to promote compartmentalized translation initiation of the HIV-1 genomic RNA. Nucleic Acids Res, 41, 6286–99.

5. Soto-Rifo, R., Rubilar, P.S., Limousin, T., de Breyne, S., Décimo, D. and Ohlmann, T. (2012) DEAD-box protein DDX3 associates with eIF4F to promote translation of selected mRNAs. Embo J, 31, 3745–56.

6. Wilkins, K.C., Schroeder, T., Gu, S., Revalde, J.L. and Floor, S.N. (2024) A novel reporter for helicase activity in translation uncovers DDX3X interactions. RNA, 30, 1041–1057.

7. Ryan, C.S. and Schröder, M. (2022) The human DEAD-box helicase DDX3X as a regulator of mRNA translation. Front. Cell Dev. Biol., 10, 1033684.

8. Lennox, A.L., Hoye, M.L., Jiang, R., Johnson-Kerner, B.L., Suit, L.A., Venkataramanan, S., Sheehan, C.J., Alsina, F.C., Fregeau, B., Aldinger, K.A., et al. (2020) Pathogenic DDX3X Mutations Impair RNA Metabolism and Neurogenesis during Fetal Cortical Development. Neuron, 106, 404–420.e8.

9. Valentin-Vega, Y.A., Wang, Y.-D., Parker, M., Patmore, D.M., Kanagaraj, A., Moore, J., Rusch, M., Finkelstein, D., Ellison, D.W., Gilbertson, R.J., et al. (2016) Cancer-associated DDX3X mutations drive stress granule assembly and impair global translation. Sci. Rep., 6, 25996.

10. Vellky, J.E., McSweeney, S.T., Ricke, E.A. and Ricke, W.A. (2020) RNA-binding protein DDX3 mediates posttranscriptional regulation of androgen receptor: A mechanism of castration resistance. Proc. Natl. Acad. Sci., 117, 28092–28101.

11. Park, J.T. and Oh, S. (2022) The translational landscape as regulated by the RNA helicase DDX3. BMB Rep., 55, 125–135.

12. Lai, M.C., Wang, S.W., Cheng, L., Tarn, W.Y., Tsai, S.J. and Sun, H.S. (2013) Human DDX3 interacts with the HIV-1 Tat protein to facilitate viral mRNA translation. PLoS One, 8, e68665.

13. Schmidt, N., Lareau, C.A., Keshishian, H., Ganskih, S., Schneider, C., Hennig, T., Melanson, R., Werner, S., Wei, Y., Zimmer, M., et al. (2021) The SARS-CoV-2 RNA–protein interactome in infected human cells. Nat. Microbiol., 6, 339–353.

14. Kukhanova, M.K., Karpenko, I.L. and Ivanov, A.V. (2020) DEAD-box RNA Helicase DDX3: Functional Properties and Development of DDX3 Inhibitors as Antiviral and Anticancer Drugs. Mol. Basel Switz., 25, 1015.

15. Gu, L., Fullam, A., McCormack, N., Höhn, Y. and Schröder, M. (2017) DDX3 directly regulates TRAF3 ubiquitination and acts as a scaffold to co-ordinate assembly of signalling complexes downstream from MAVS. Biochem. J., 474, 571–587.

16. Gu, L., Fullam, A., Brennan, R. and Schröder, M. (2013) Human DEAD Box Helicase 3 Couples IκB Kinase ε to Interferon Regulatory Factor 3 Activation. Mol. Cell. Biol., 33, 2004–2015.

17. Schröder, M., Baran, M. and Bowie, A.G. (2008) Viral targeting of DEAD box protein 3 reveals its role in TBK1/IKKepsilon-mediated IRF activation. Embo J, 27, 2147–57.

18. Anastasakis, D.G., Jacob, A., Konstantinidou, P., Meguro, K., Claypool, D., Cekan, P., Haase, A.D. and Hafner, M. (2021) A non-radioactive, improved PAR-CLIP and small RNA cDNA library preparation protocol. Nucleic Acids Res., 49, 10.

19. Polash, A.H. and Hafner, M. (2026) PCLIPtools: a robust framework for identifying RNA-protein interaction sites from PAR-CLIP experiments. Nucleic Acids Res., 54, gkag062.

20. Martin, M. (2011) Cutadapt removes adapter sequences from high-throughput sequencing reads. EMBnet.journal, 17, 10–12.

21. Dobin, A., Davis, C.A., Schlesinger, F., Drenkow, J., Zaleski, C., Jha, S., Batut, P., Chaisson, M. and Gingeras, T.R. (2013) STAR: ultrafast universal RNA-seq aligner. Bioinforma. Oxf. Engl., 29, 15–21.

22. Li, H., Handsaker, B., Wysoker, A., Fennell, T., Ruan, J., Homer, N., Marth, G., Abecasis, G., Durbin, R., and 1000 Genome Project Data Processing Subgroup (2009) The Sequence Alignment/Map format and SAMtools. Bioinformatics, 25, 2078–2079.

23. Quinlan, A.R. and Hall, I.M. (2010) BEDTools: a flexible suite of utilities for comparing genomic features. Bioinformatics, 26, 841–842.

24. Dyer, S.C., Austine-Orimoloye, O., Azov, A.G., Barba, M., Barnes, I., Barrera-Enriquez, V.P., Becker, A., Bennett, R., Beracochea, M., Berry, A., et al. (2025) Ensembl 2025. Nucleic Acids Res., 53, D948–D957.

25. Liao, Y., Smyth, G.K. and Shi, W. (2014) featureCounts: an efficient general purpose program for assigning sequence reads to genomic features. Bioinformatics, 30, 923–930.

26. Perez, G., Barber, G.P., Benet-Pages, A., Casper, J., Clawson, H., Diekhans, M., Fischer, C., Gonzalez, J.N., Hinrichs, A.S., Lee, C.M., et al. (2025) The UCSC Genome Browser database: 2025 update. Nucleic Acids Res., 53, D1243–D1249.

27. Patro, R., Duggal, G., Love, M.I., Irizarry, R.A. and Kingsford, C. (2017) Salmon provides fast and bias-aware quantification of transcript expression. Nat. Methods, 14, 417–419.

28. Pertea, G. and Pertea, M. (2020) GFF Utilities: GffRead and GffCompare. F1000Research, 9, 304.

29. Muys, B.R., Anastasakis, D.G., Claypool, D., Pongor, L., Li, X.L., Grammatikakis, I., Liu, M., Wang, X., Prasanth, K.V., Aladjem, M.I., et al. (2021) The p53-induced RNA-binding protein ZMAT3 is a splicing regulator that inhibits the splicing of oncogenic CD44 variants in colorectal carcinoma. Genes Dev., 35, 102–116.

30. Lorenz, R., Bernhart, S.H., Höner zu Siederdissen, C., Tafer, H., Flamm, C., Stadler, P.F. and Hofacker, I.L. (2011) ViennaRNA Package 2.0. Algorithms Mol. Biol., 6, 26.

31. Robinson, J.T., Thorvaldsdóttir, H., Winckler, W., Guttman, M., Lander, E.S., Getz, G. and Mesirov, J.P. (2011) Integrative genomics viewer. Nat. Biotechnol., 29, 24–26.

32. Szklarczyk, D., Kirsch, R., Koutrouli, M., Nastou, K., Mehryary, F., Hachilif, R., Gable, A.L., Fang, T., Doncheva, N.T., Pyysalo, S., et al. (2023) The STRING database in 2023: protein-protein association networks and functional enrichment analyses for any sequenced genome of interest. Nucleic Acids Res., 51, D638–D646.

33. R Core Team (2021) R: a language and environment for statistical computing R Foundation for Statistical Computing, Vienna, Austria.

34. Love, M.I., Huber, W. and Anders, S. (2014) Moderated estimation of fold change and dispersion for RNA-seq data with DESeq2.

35. Kolde, R. (2019) pheatmap: Pretty heatmaps.

36. Wickham, H. (2016) ggplot2: Elegant graphics for data analysis Springer-Verlag New York.

37. Sánchez-Aparicio, M.T., Garcin, D., Rice, C.M., Kolakofsky, D., García-Sastre, A. and Baum, A. (2017) Loss of Sendai virus C protein leads to accumulation of RIG-I immunostimulatory defective interfering RNA. J. Gen. Virol., 98, 1282–1293.

38. Oh, S., Flynn, R.A., Floor, S.N., Purzner, J., Martin, L., Do, B.T., Schubert, S., Vaka, D., Morrissy, S., Li, Y., et al. (2016) Medulloblastoma-associated DDX3 variant selectively alters the translational response to stress. Oncotarget, 7, 28169–28182.

39. Gong, C., Krupka, J.A., Gao, J., Grigoropoulos, N.F., Giotopoulos, G., Asby, R., Screen, M., Usheva, Z., Cucco, F., Barrans, S., et al. (2021) Sequential inverse dysregulation of the RNA helicases DDX3X and DDX3Y facilitates MYC-driven lymphomagenesis. Mol. Cell, 81, 4059–4075.e11.

40. Trotta, E. (2014) On the Normalization of the Minimum Free Energy of RNAs by Sequence Length. PLoS ONE, 9, e113380.

41. Kato, A., Cortese-Grogan, C., Moyer, S.A., Sugahara, F., Sakaguchi, T., Kubota, T., Otsuki, N., Kohase, M., Tashiro, M. and Nagai, Y. (2004) Characterization of the Amino Acid Residues of Sendai Virus C Protein That Are Critically Involved in Its Interferon Antagonism and RNA Synthesis Down-Regulation. J. Virol., 78, 7443–7454.

42. Curran, J., Latorre, P. and Kolakofsky, D. (1998) Translational Gymnastics on the Sendai Virus P/C mRNA. Semin. Virol., 8, 351–357.

43. Mizuguchi, H., Nakanishi, T., Kondoh, M., Nakagawa, T., Nakanishi, M., Matsuyama, T., Tsutsumi, Y., Nakagawa, S. and Mayumi, T. (1999) Fusion of Sendai virus with liposome depends on only F protein, but not HN protein. Virus Res., 59, 191–201.

44. Faísca, P. and Desmecht, D. (2007) Sendai virus, the mouse parainfluenza type 1: A longstanding pathogen that remains up-to-date. Res. Vet. Sci., 82, 115–125.

45. Yoshizumi, T., Imamura, H., Taku, T., Kuroki, T., Kawaguchi, A., Ishikawa, K., Nakada, K. and Koshiba, T. (2017) RLR-mediated antiviral innate immunity requires oxidative phosphorylation activity. Sci. Rep., 7, 5379.

46. Cheng, M., Niu, Y., Fan, J., Chi, X., Liu, X. and Yang, W. (2018) Interferon down-regulation of miR-1225-3p as an antiviral mechanism through modulating Grb2-associated binding protein 3 expression. J. Biol. Chem., 293, 5975–5986.

47. Herdy, B., Mayer, C., Varshney, D., Marsico, G., Murat, P., Taylor, C., D’Santos, C., Tannahill, D. and Balasubramanian, S. (2018) Analysis of NRAS RNA G-quadruplex binding proteins reveals DDX3X as a novel interactor of cellular G-quadruplex containing transcripts. Nucleic Acids Res., 46, 11592–11604.

48. Toyama, Y., Takeuchi, K. and Shimada, I. (2025) Regulatory role of the N-terminal intrinsically disordered region of the DEAD-box RNA helicase DDX3X in selective RNA recognition. Nat. Commun., 16, 7762.

49. Varshney, D., Cuesta, S.M., Herdy, B., Abdullah, U.B., Tannahill, D. and Balasubramanian, S. (2021) RNA G-quadruplex structures control ribosomal protein production. Sci. Rep., 11, 22735.

50. Brown, N.P., Vergara, A.M., Whelan, A.B., Guerra, P. and Bolger, T.A. (2021) Medulloblastoma-associated mutations in the DEAD-box RNA helicase DDX3X/DED1 cause specific defects in translation. J. Biol. Chem., 296, 100296.

51. Irie, T., Okamoto, I., Yoshida, A., Nagai, Y. and Sakaguchi, T. (2014) Sendai Virus C Proteins Regulate Viral Genome and Antigenome Synthesis To Dictate the Negative Genome Polarity. J. Virol., 88, 690–698.

52. Chen, H., Charles, P.D., Gu, Q., Liberatori, S., Robertson, D.L., Palmarini, M., Wilson, S.J., Mohammed, S. and Castello, A. (2025) Omics Analyses Uncover Host Networks Defining Virus-Permissive and -Hostile Cellular States. Mol. Cell. Proteomics, 24.

53. Bedsaul, J.R., Zaritsky, L.A. and Zoon, K.C. (2016) Type I Interferon-Mediated Induction of Antiviral Genes and Proteins Fails to Protect Cells from the Cytopathic Effects of Sendai Virus Infection. J. Interferon Cytokine Res., 36, 652–665.

