## Supplementary Figures for "The human DEAD-box protein DDX3X regulates host and viral mRNA translation during Sendai Virus infection"

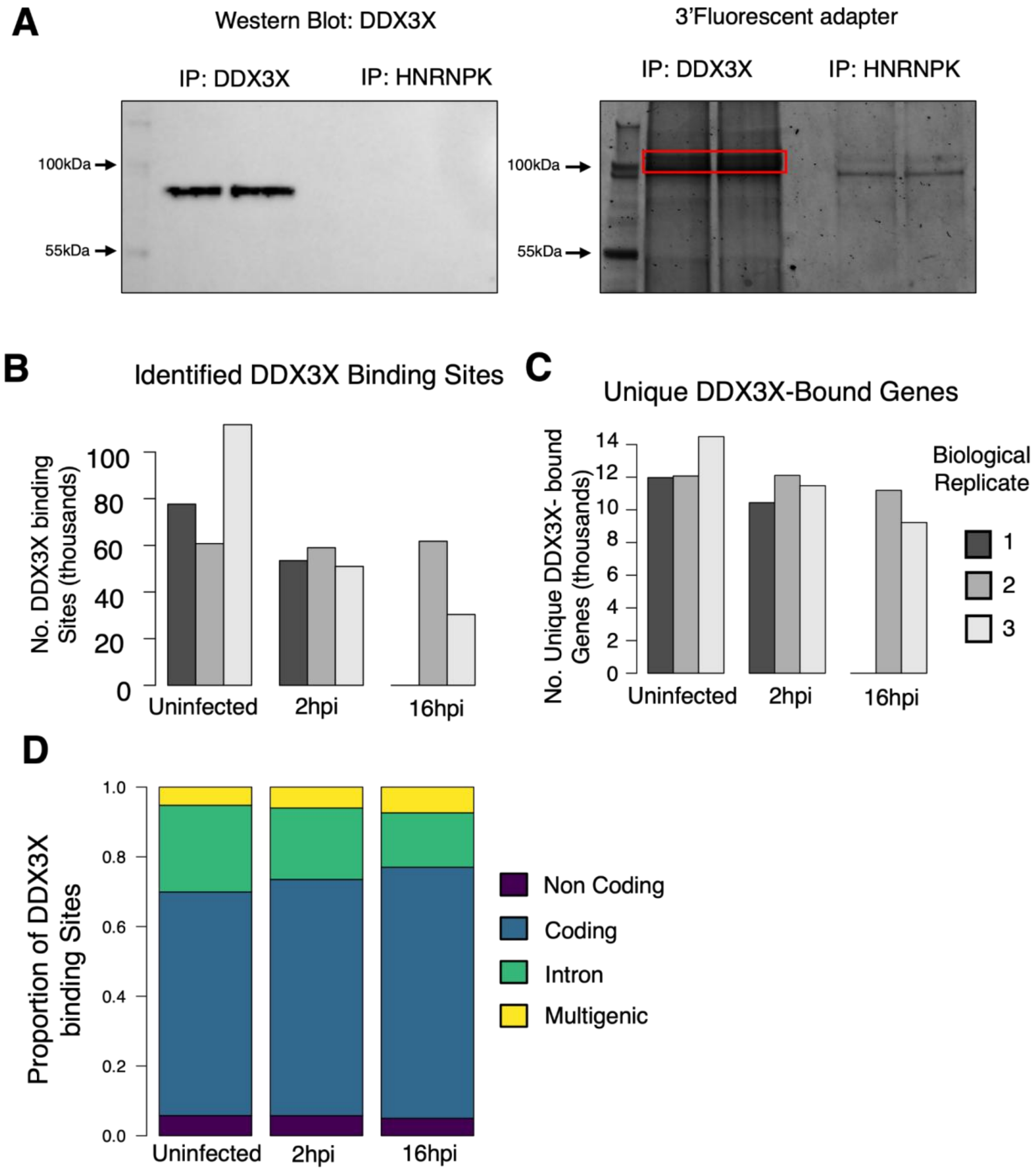

**Figure S1: PAR-CLIP of DDX3X in SeV-infected and uninfected HEK293T cells.** (A) Left: Western blot against DDX3X of PAR-CLIP samples using either anti-DDX3X or anti-HNRNPK (for comparison) for IP. Right: Scan for Alexa-Fluor 647 fluorescence of SDS-PAGE gel of the same samples. The region corresponding to resolved DDX3X-mRNA complexes excised from the gel for library preparation is highlighted. (B) Per-sample number of unique DDX3X binding sites identified from clusters of DDX3X-crosslinked reads bearing T-to-C conversions (C) Count of unique genes identified from annotated DDX3X binding sites. Individual bars at each timepoint refer to individual biological replicates (Uninfected and 2hr n = 3, 8hr and 16hr n = 2). hpi = hours-post Sendai Virus infection. (D) Stacked bar plot comparing proportions of DDX3X-crosslinked reads mapping to protein-coding exon, protein-coding intron, non-coding, and multigenic regions of the genome in uninfected and infected cells (using the mean count of reads per sample).

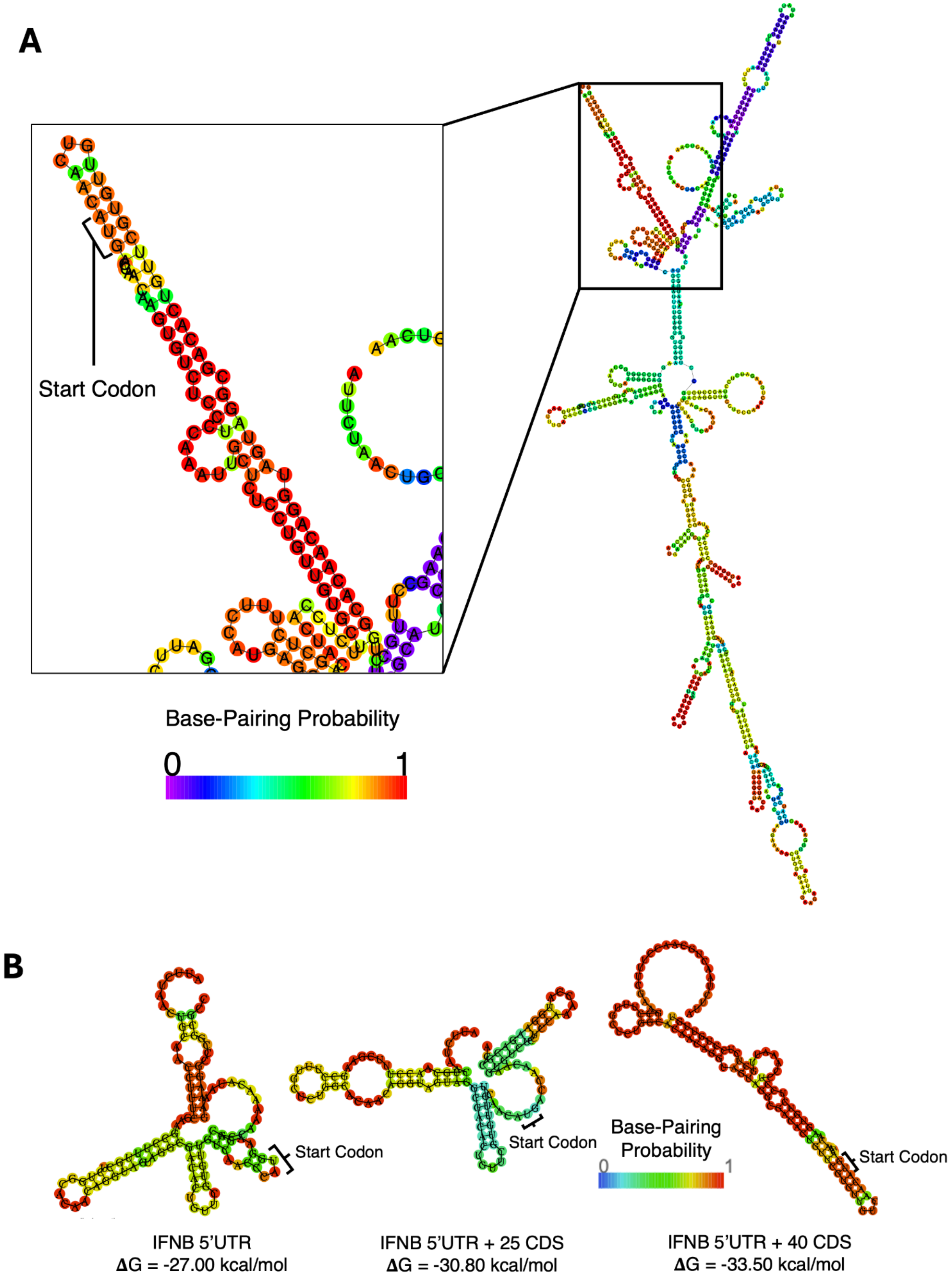
A
Start Codon
Base-Pairing Probability
0
1
B
Start Codon
Start Codon
Base-Pairing Probability
0
1
Start Codon
IFNB 5'UTR
ΔG = -27.00 kcal/mol
IFNB 5'UTR + 25 CDS
ΔG = -30.80 kcal/mol
IFNB 5'UTR + 40 CDS
ΔG = -33.50 kcal/mol

**Figure S2:** (A) Minimum free energy secondary structures of IFNB1 mRNA predicted by Vienna RNAfold (30). Colours of paired/unpaired residues correspond to probabilities of being paired/unpaired, respectively. (B) Predicted MFE structures of IFNB reporters predicted by Vienna RNAfold (30). Each folded sequence is of equal length to the longest reporter insert (IFNB 5'UTR + 40 CDS, 113nt), with the shorter inserts including portions of the downstream FLUC CDS.

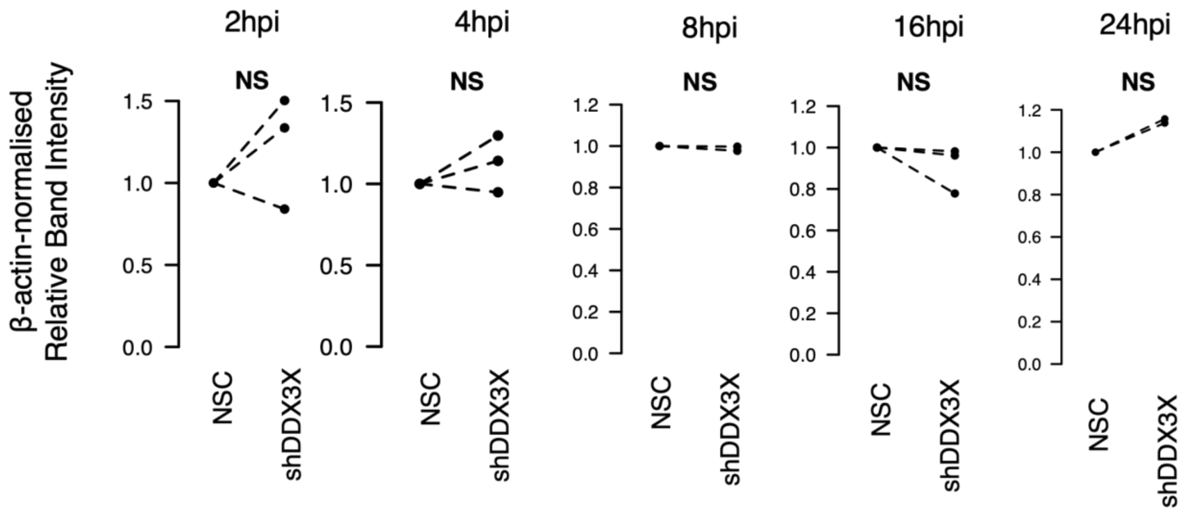


**Figure S3: Quantification of effects of DDX3X knockdown on SeV N levels.** Comparison of intensities of western blot SeV N bands normalised to corresponding β-actin bands between NSC and shDDX3X cells after 72 h doxycycline treatment (n = 2 for 8, and 24 hpi, n = 3 for 2, 4, and 16 hpi).

**Figure S4: Diversity of structural features occurring proximal to the start codon.** A random sample of 10000 sequences from DDX3X-bound transcripts, consisting of the transcript AUG ± 50nt, were folded in silico and features of these structures were analysed. (A) Hierarchically-clustered heatmap of base-pairing nucleotide residues within the MFE structures. Base-paired nucleotides are coloured purple and orange, and non-base paired nucleotides are white. (B) Hierarchically-clustered heatmap of number of enclosing base pairs in predicted MFE structures, where higher values (and darker colour) indicate nucleotides positioned within longer stretches of base-pairing.

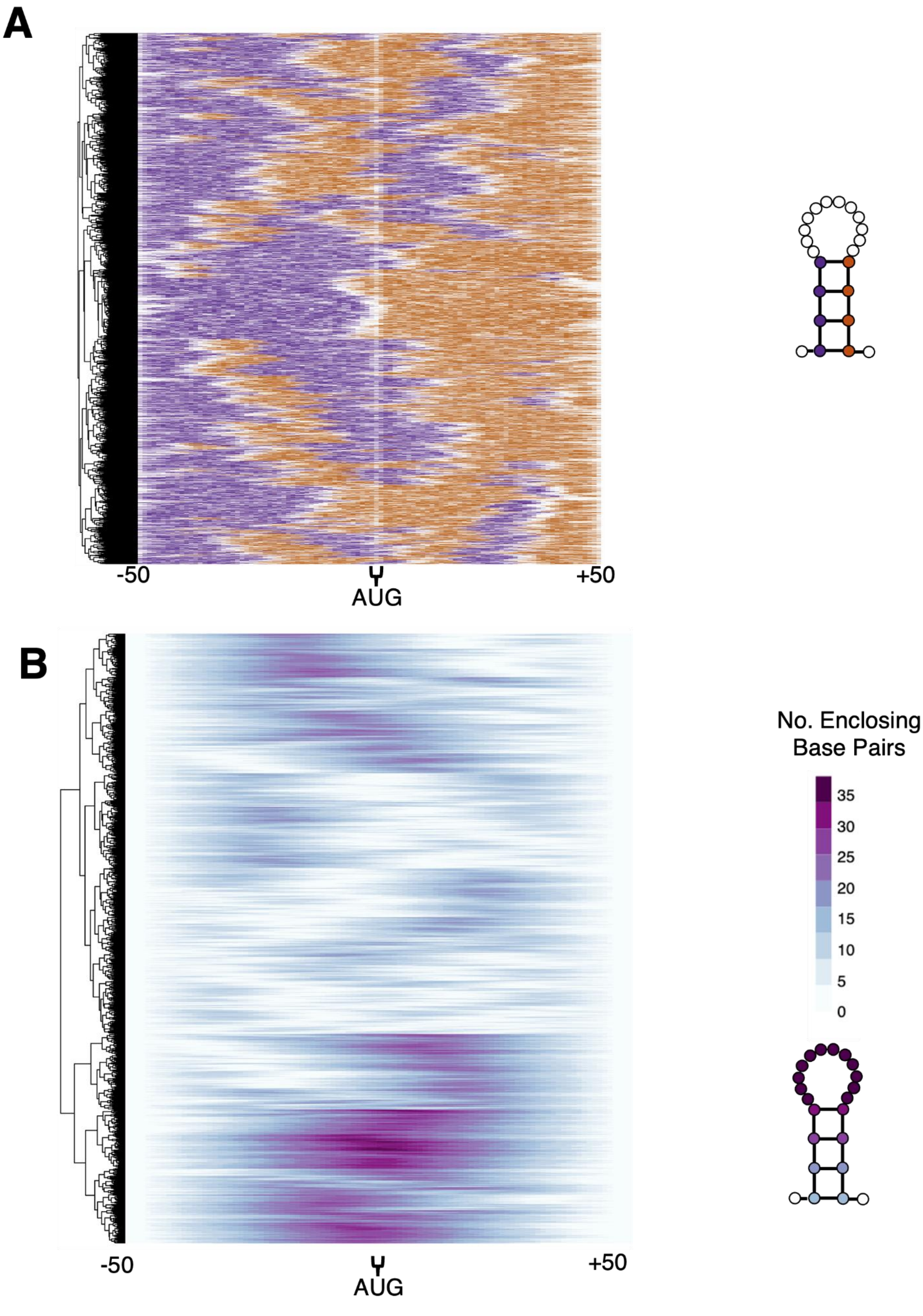
A
-50
AUG
+50
B
No. Enclosing Base Pairs
35
30
25
20
15
10
5
0
-50
AUG
+50

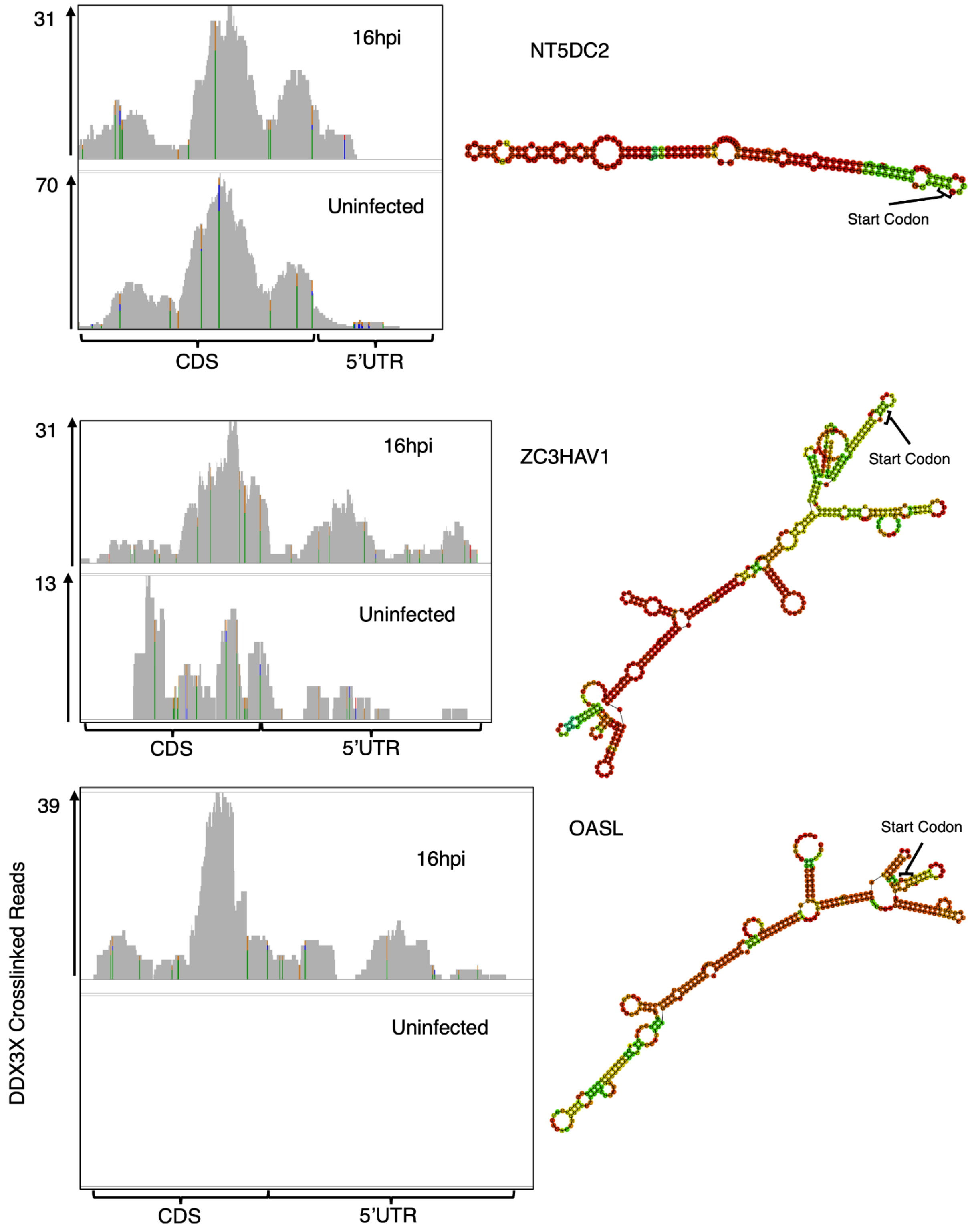


**Figure S5: Examples of DDX3X targets which display binding spanning the 5′UTR and CDS proximal to sites of predicted secondary structures.** Left: IGV screenshots showing DDX3Xcrosslinked PAR-CLIP reads aligned to target genes. PAR-CLIP T-to-C conversions are indicated as A-to-G mis-matches. Colored sites possess a large proportion of reads with nucleotide mismatches, with the ratio of A,T,C,G shown by the ratio of green, red, blue and orange, respectively.
